# Adolescent morphine exposure impairs cognitive flexibility in female but not male mice

**DOI:** 10.1101/2023.04.09.536177

**Authors:** Aditii Wakhlu, Annabel Engelhardt, Eden M. Anderson, Elaine Grafelman, Abbigail Ouimet, Matthew C. Hearing

## Abstract

Use of prescription opioids continues to rise, especially in adolescent individuals. As adolescence is a critical development window for higher order cognitive functions, thus opioid exposure during this period may have significant long-lasting effects on cognitive function and predisposition individuals to be at greater risk of developing opioid use later in life. Here, we examine previously explored effects of opioid exposure during adolescence on affect-related behavior, motivation, and cognitive flexibility. We find that a two-week exposure to non-contingent morphine during adolescence (i.e., post-weaning) does not alter performance in an elevated plus maze, forced swim test, or motivation for appetitive reward in male or female mice when tested during adolescence or adulthood. Examination of how adolescent morphine impacts cognition revealed impairments in visual-based discriminative learning and cognitive flexibility in female but not male mice, as assessed using an operant-based attentional set-shifting task. Unexpectedly, deficits in discriminative learning are observed when testing occurred during adolescence but not adulthood, whereas impaired performance in the extradimensional shift remained impaired into adulthood. The data indicate that opioid exposure during adolescence has a greater impact on cognitive function in female mice and that these deficits may be more widespread during acute withdrawal periods, while deficits in flexibility more enduring.

## Introduction

Use of opioid-based drugs carry a risk of dependence and misuse. While use of prescription opioids has continued to increase in all populations in recent years, adolescent use has dramatically increased, with a higher prevalence of use in adolescent females compared to males (Hudgins JD et al., 2019;Hudgins JD et al., 2019). In humans, development of opioid use disorders (OUD) is associated with a reduction in the ability to flexibly regulate behaviors (Anderson EM et al., 2021;Goldstein RZ and Volkow ND, 2011;Lee TM et al., 2005;Li CS and Sinha R, 2008;Zeng H et al., 2013). Cognitive flexibility undergoes changes across childhood and adolescence and represents a critical transitional period for normal development of executive control over behavior (Cragg L and Chevalier N, 2012;Dajani DR and Uddin LQ, 2015). As exposure to prescription opioids in adolescence is highly predictive of subsequent opioid misuse and development of OUD (Miech R et al., 2015), adolescence may also mark a period of unique vulnerability to the effects of opioids on cognitive control.

While deficits in cognitive control are a hallmark of OUD, it is unclear whether this phenotype is the result of chronic opioid exposure, or rather is a risk factor predating the development of OUDs. For example, innate individual differences in cue reactivity, incentive motivation, and cognitive flexibility can predict subsequent drug use in humans and in preclinical animal models (Flagel et al., 2009; Saunders and Robinson, 2013; Shnitko et al., 2019). Further, age of exposure (Meich 2015, Chen et al., 2009; Barker et al., 2017) and biological sex is known to influence the propensity to develop substance use disorders and susceptibility to drug behavior (Anderson EM,Engelhardt A,Demis S,Porath E and Hearing MC, 2021;Barker JM et al., 2017;Becker JB et al., 2012;Cicero TJ et al., 2003;Kennedy AP et al., 2013;Lynch WJ and Carroll ME, 1999)(Serdarevic et al., 2017; Kennedy et al., 2013). Recent work indicates that sex and the developmental age of exposure to drugs such as alcohol interact to influence susceptibility on developing impairments in cognitive control (Barker et al., 2017). While biological sex has been shown to influence susceptibility to opioid-induced impairments in cognitive flexibility, whether interactions between age and sex occurs with opioid exposure remains unknown.

The prolonged developmental trajectory of the prefrontal cortex (PFC), necessary for acquisition of executive function and complex cognitive abilities, may increase susceptibility to disruption of normal functioning during adolescence. Cognitive flexibility, a critical component of executive function, undergoes changes across childhood and adolescence and the critical transitional period is for normal development of executive control (Cragg & Chevalier, 2012, Dajani and Uddin, 2015). Deficits in cognitive flexibility remain one of the most consistently documented cognitive problems associated with substance use disorders. Addiction vulnerability to and severity can be exacerbated by deficits in cognitive flexibility by increasing drug-seeking behaviors, relapse, and impairment in the ability to resist habitual drug use. Past studies in the lab have shown that exposure to remifentanil leads to lasting changes in the PFC hypoactive basal state evidenced by a decrease in ex vivo excitability that is paralleled by an increase in firing capacity of layer 5/6 pyramidal neurons in the prelimbic - which aligns with deficits in cognitive flexibility as assessed using an operant-based attentional set shifting task.

## Materials and Methods

### Subjects

Adult male and female wild-type C57BL/6 mice were bred in-house or commercially purchased (Jackson Laboratory) and maintained in a temperature and humidity-controlled room. Animal use was approved by the Institutional Animal Care and Use Committee at Marquette University.

### Morphine exposure

Twenty-four hours following weaning (postnatal day (PND) 22-29), mice received 14 days of noncontingent subcutaneous morphine (10 mg/kg) or saline injections (excluding weekends). Successful delivery of morphine was denoted 30 minutes following exposure based on morphine-induced straub tail response (Nath C et al., 1994). To test the effects of adolescent morphine exposure on behavioral measures during adolescence, mice began behavioral testing 24-h following the final injection (PND35-39). To determine if any observed effects of adolescent exposure persisted into adulthood, behavioral testing began 21-24-d following the final injection.

### Behavioral assessments

A timeline of behavioral assessments is provided in Figure 1A based on age of testing post injection. Testing in the **elevated plus maze** (EPM; San Diego Instruments) occurred for 5 min as previously described (Anderson et al., 2019) in the morning on day one of testing. Percent time in the open arms was calculated as total time in the open arms divided by total time in the maze. Approximately 8-h later, testing for immobility in the **forced swim test** (FST) was done in a beaker filled with 25 ± 2ºC tap water and allowed to habituate for 2 minutes followed by 4 minutes of testing during which the time spent immobile and the number of immobile episodes were tracked using a side-mounted camera and AnyMaze software with immobility sensitivity of 85% and a minimum immobility episode time of 250 milliseconds.as described (Anderson et al., 2019). Following the FST, mice were food deprived for 16-24-h and began food training in the **attention set-shifting task** (ASST) to measure cognitive flexibility as previously described (Anderson EM et al., 2021;Anderson EM,Engelhardt A,Demis S,Porath E and Hearing MC, 2021;Anderson EM et al., 2021). During *food training*, a fixed ratio 1 schedule was used, whereby a right lever response (opposite lever to the reinforced lever during self - administration) resulted in delivery of a 50% liquid Ensure® reward. Food training sessions were 30 minutes in length and required mice to earn at least 50 rewards to progress to lever training. During *lever training*, levers were extended pseudorandomly with no more than two consecutive extensions of each lever. Mice were required to press the lever within 10 seconds of extension to receive a 50% liquid Ensure® reward, absence of which was deemed as an omission. Lever training consisted of a total of 90 trials (45 of each the left and right lever), with each followed by a 20 second time out. Mice had to reach criterion of five or fewer omissions on two consecutive days. Once lever training criterion was reached, *lever bias* was assessed through 7 trials, during which both the left and the right lever were reinforced. The following day, mice underwent visual cue testing until 150 trials or 10 consecutive correct responses were made. During the visual cue test, the correct response was on the lever below the illuminated cue light. Incorrect responses resulted in only the time out (no reward). Failure to respond (i.e. omission) was tracked but not counted towards a trial to criterion. The day following visual cue testing, *extradimensional shift testing* was conducted. The reinforced lever was always the lever opposite of the lever bias; however, the cue light was presented similar to that during the visual cue test. The day after criterion of the extradimensional shift was reached (i.e. 10 consecutive correct responses), reversal testing was conducted with the reinforced lever always that of the bias and the cue light presented identical to that during the visual cue test.

**Figure 1:**
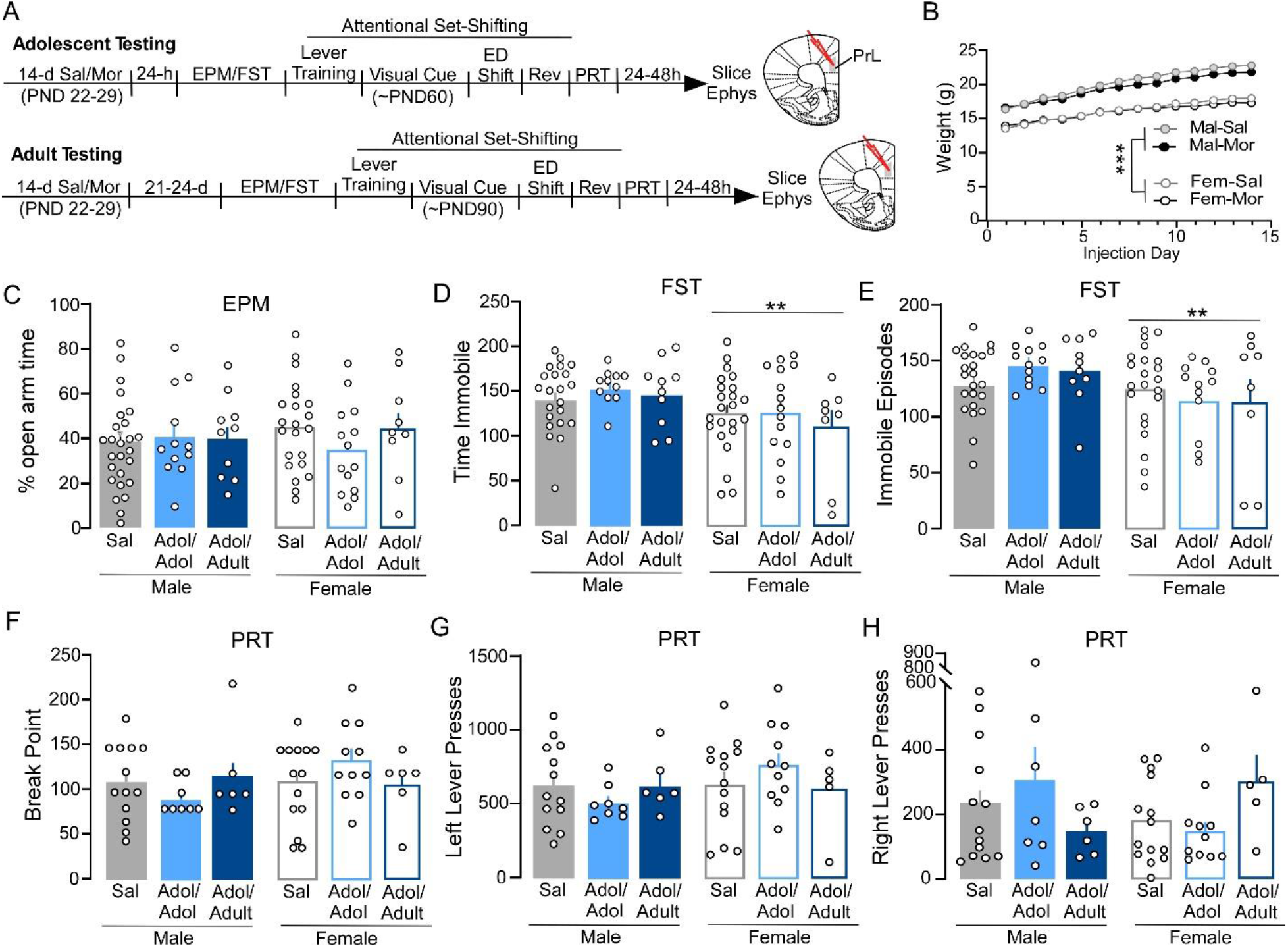
**A)** Timeline of withdrawal and behavioral assessments following 14-d of non-contingent morphine exposure in male and female mice. **B)** Body weights of male and female mice taken daily prior to injections. **C)** Percent time spent in the open arm of an elevated plus maze (EPM) in male (left, filled) and female (right, unfilled) mice treated with saline (gray) or morphine with subsequent testing during adolescence (adol/adol, light blue) and adulthood (adol/adult, dark blue) following saline (gray). **D)** Time spent immobile during a forced swim test (FST). **E)** Number of immobile episodes during an FST. **F)** Break points, **G)** total left lever presses, and **H**, total right lever presses during a progressive ration test (PRT) of motivation for an appetitive reward. **p<0.01 females versus males.

For further behavioral assessment, the types of errors were analyzed based on previously published methods(Brady AM and Floresco SB, 2015). Tests were divided into blocks of 16 trials, not included trials were there was an omitted responses. For visual cue testing, *initial errors* were errors that were made within each block until there were less than six in a single block. Once there were less than six errors in a single block, errors in all subsequent blocks were characterized as *regressive errors*. For the extradimensional set shift test, tests were divided into bins of 16 trials. Errors that were made based on the previous rule used with the visual cue test such that the response was made on the lever under the illuminated cue light were tagged and were considered *perseverative* until less than six errors in a single bin were made. Errors in the next bin and subsequent bins were considered regressive errors. Never reinforced errors were those that were incorrect but the response was not on the lever underneath the illuminated cue light. For set-shifting experiments, data was excluded if >15 omissions in a given test was recorded as this may reflect mechanical issues or motoric issues. If these omissions were evident only during the reversal test, visual cue and extradimensional data was still used. Conversely, if these omissions were during the visual cue or extradimensional test, then all data was excluded as responding during these tests may have influenced responding during the extradimensional test or reversal test, respectively. After conclusion of the reversal test, mice underwent a **progressive ratio test** (PRT) to assess motivation using the same liquid reward. Responses on the left lever were reinforced and the responses required to obtain the next reward progressively increased with each reward obtained ((5e^.2*n^)-5; (Richardson NR and Roberts DC, 1996). Testing continued until 30min elapsed without a response or a total of 1.5 hours. It should be noted that a majority of mice were still responding at the end of the test session and is a potential limitation to the interpretation of these findings.

### Ex-vivo slice electrophysiology

Whole-cell recordings were performed in layer 5/6 pyramidal neurons in the prelimbic cortex as previously described (Hearing M et al., 2013). Recordings were performed in mice undergoing behavioral testing and mice naïve of testing. Sutter Integrated Patch Amplifier (IPA) with Igor Pro (Wave Metrics, Inc.) was used for the data acquisition software. Rheobase and action potential frequency were assessed using a 20 pA 1-second current-step injection from 0-500 pA. Duration of the action potential was quantified as the time it took to reach half the amplitude of the initial action potential. Afterhyperpolarization (AHP) amplitude was measured from the action potential threshold to the maximum afterhyperpolarization amplitude of the initial action potential.

### Data analysis

For analysis of weights during injection, a 2-way RM ANOVA was run with a 4 group (male-saline, female-saline, male-morphine, female-morphine) x day (1-14) comparison. For experiments where saline groups were combined, a 2 (male, female) x 3 (saline, adolescent exposed – adolescent tested, and adolescent exposed/adult tested), 30d remifentanil) ANOVA was used. With the exception of current-spike analysis for ex vivo recordings, 2 (male, female) x 2 (saline, morphine) ANOVAs were used to test for statistical significance for initial rheobase studies, elevated plus maze, forced swim test, and attention set-shifting studies. Current-spike analysis was analyzed using a 2 (saline, morphine) way repeated-measures ANOVA. Rheobase comparing only two groups (e.g. males versus females) was analyzed using an independent-samples t-test. For experiments where saline groups were combined a 2 (male, female) x 3 (saline, adolescent exposed/adolescent, adolescent exposed/adult tested) ANOVA. Prior to combining saline groups, statistical comparisons were performed using a student t-test to ensure that no significant differences were observed across adolescent and adult controls (**Table 1**.) When interactions were found, main effects were not reported. Student-Newman-Keuls post-hoc comparisons were conducted when necessary. IPSCs/EPSCs were analyzed using MiniAnalysis software using a 5pA detection threshold (Synaptosoft). Data was analyzed using Sigma Plot (Systat Software, Inc) and graphed using Prism 7.0 (GraphPad). The threshold for statistical significance was p<0.05. Shapiro-Wilk test of normality and an equal variance assumption was used, when applicable. Statistical outliers were defined as being at least 2 standard deviations away from the mean and were removed from analyses. Across all treatment groups (8 total), 6 outliers were identified with no more than 1 per group removed. For electrophysiology studies, no more than 3 recordings were used for a given metric (e.g., mEPSCs) from the same animal to reduce data bias.

**Table 1.** Comparison of behavioral measures taken from saline-treated male and female mice show no difference in mice when tested during adolescence versus adulthood.

| Behavior Test/Endpoint | Female (Saline) |  |  |  |  | Male (Saline) |  |  |  |  |
| --- | --- | --- | --- | --- | --- | --- | --- | --- | --- | --- |
|  | Adolescent |  | Adult |  | t (df), p value | Adolescent |  | Adult |  | t (df), p value |
| | N | mean ± sem | N | mean ± sem | | N | mean ± sem | N | mean ± sem | |
| EPM (% open arm time) | 15 | 43.3 ± 5.2 | 7 | 50.1 ± 7.4 | t(20)= -0.75, p=0.46 | 15 | 33.3 ± 4.4 | 11 | 42.98 ± 6.9 | t(24)=-1.23, p=0.23 |
| FST (Time Immobile) | 16 | 120 ± 7.6 | 7 | 107 ± 22.7 | t(21)= 0.71, p=0.48 | 12 | 131.0 ± 11.3 | 11 | 141.2 ± 6.8 | t(21)=-0.75, p=0.46 |
| FST (Immobile Episodes) | 16 | 124 ± 6.9 | 7 | 117 ± 22.4 | t(21)= 0.39, p=0.69 | 12 | 122.1 ± 8.1 | 11 | 138.9 ± 9.2 | t(21)=-1.38, p=0.18 |
| PRT (Left Lever Presses) | 8 | 662 ± 102 | 6 | 558 ± 140 | t(12)= 0.55, p=0.59 | 7 | 709 ± 83.2 | 6 | 532 ± 133 | t(11)=1.16, p=0.27 |
| PRT (Right Lever Presses) | 8 | 154 ± 38.2 | 6 | 205 ± 64.1 | t(12)= -0.71, p=0.49 | 7 | 309 ± 82.9 | 6 | 136 ± 32.2 | t(11)=1.82, p=0.51 |
| PRT (Break Point) | 8 | 113 ± 16.1 | 6 | 97.3 ± 22.2 | t(12)= 0.61, p=0.56 | 7 | 121 ± 11.5 | 6 | 90.5 ± 21.1 | t(11)=1.31, p=0.21 |
| ASST (Lever Training, days) | 8 | 7.25 ± 0.53 | 7 | 8.14 ± 1.42 | t(13)= -0.62, p=0.55 | 7 | 6.4 ± 0.84 | 6 | 6.7 ± 1.05 | t(11)=-0.18, p=0.86 |
| ASST (Vis Cue, Trials) | 8 | 79.6 ± 14.6 | 7 | 72.9 ± 11.9 | t(13)= 0.35, p=0.73 | 7 | 75 ± 9.3 | 6 | 87.3 ± 21.6 | t(11)=-0.55, p=0.59 |
| ASST (ED Shift, Trials) | 8 | 76.5 ± 8.4 | 7 | 72.0 ± 5.8 | t(13)= 0.43, p=0.67 | 7 | 94.1 ± 11.9 | 6 | 75.5 ± 9.9 | t(11)= 1.17, p=0.27 |
| ASST (Reversal, Trials) | 8 | 125 ± 13.7 | 7 | 117 ± 13.8 | t(13)= 0.41, p=0.69 | 7 | 114.1 ± 22.2 | 6 | 131.2 ± 33.3 | t(11)= -0.44, p=0.67 |

## Results

### Effect of adolescent morphine exposure on performance in EPM, FST, and PRT during adolescence or adulthood

Twenty-four hours after weaning, mice received 14-daily injections of morphine (10 mg/kg; sc) at postnatal day (PND) 22-29. Individual animal weights were taken prior to daily injections. Comparison of weights done without consideration for behavioral testing timepoint showed an interaction between group (male-saline, female-saline, male-morphine, female-morphine; F_(3,1007)_= 67.35, p<0.001) and day (F_(39,1007)_ = 4.7, p<0.001), with male mice treated with morphine or saline exhibiting greater weights than female counterparts. However no significant effect of treatment within sex was observed, suggesting that the morphine regimen did not produce physical dependence (Figure 1B).

Following morphine exposure, mice were randomly selected to undergo sequential behavioral testing that began during adolescence (adol/adol) or in adulthood (adol /adult) (Figure 1A). We first examined whether morphine exposure produced changes in performance in the elevated plus maze (EPM) and a forced swim test (FST). No effect of sex (male, female), treatment condition (saline, adol/adol (morphine), adol/adult (morphine)), or interaction of sex and treatment condition was observed in the percent of time spent in the EPM open arm (2-way ANOVA, sex: F_(1,92)_ = 0.2, p=0.66; condition: F_(2,92)_ = 0.26, p=0.77; interaction: F_(2,92)_=1.11, p = 0.33; Figure 1C). Alternatively, examination of coping strategies in the FST (immobility) showed that females exhibited significantly less time immobile (sex: F_(1,90)_=10.00, p=0.002; condition: F_(2,90)_=0.428, p=0.65; interaction: F_(2,90)_ = 0.464, p=0.63; Figure 1D) and fewer immobile episodes (sex: F_(1,90)_=10.22, p=0.002; condition: F_(2,90)_=0.33, p=0.72; interaction: F_(2,90)_=1.78, p=0.18) compared to their male counter counterparts regardless of treatment condition (Figure 1E).

During a progressive ratio test (PRT) of motivation for an appetitive reward, no effect of sex (F_(1,57)_=0.24, p=0.63), treatment condition (F_(2,57)_=0.39, p=0.68), or interaction of sex and treatment condition (F_(2,57)_=0.88, p=0.42) was found when comparing break point values (Figure 1F). Similarly, no differences were observed in total presses on the left (active) lever (sex: F_(1,53)_=0.574, p=0.45); condition: F_(2,53)_=0.99; interaction: (F_(2,53)_=1.08, p=0.35; Figure 1G) or right (inactive) lever (sex: F_(1,53)_=0.355, p=0.55); condition: F_(2,53)_=0.08, p=0.924; interaction: F_(2,53)_=2.73, p=0.07; Figure 1H).

### Sex-specific and age-dependent impact of adolescent morphine on cognitive flexibility

Recent findings have shown that volitional opioid administration in adult mice promotes sex-specific effects on cognitive flexibility (Anderson et al., 2021), however the impact of opioid exposure during adolescence on learning and decision-making remains unclear. To examine effects of morphine on cognitive flexibility, we used the same operant-based attentional set-shift model that resembles the Wisconsin Card Sorting Task in the sense that the stimuli are easy to detect, but rules are implicit and learned while the task is performed (Figure 2A). Morphine exposure did not alter the number of days to reach lever training criterion in male or females (sex: F_(1,61)_ = 0.11, p=0.74; condition: F_(2,61)_ = 0.15, p=0.86; interaction: F_(2,61)_ = 0.78, p=0.46). When performance was assessed during a visual cue discrimination task, adolescent morphine exposure increased the number of trials to reach criterion in female, but not male mice compared to saline counterparts when tested during adolescence. Alternatively, morphine did not alter the number of trials to criterion in either males or females compared to saline controls when tested in adulthood (*trials to criterion*, sex x treatment interaction: F_(2,61)_=6.41, p=0.003; *errors to criterion*, (F_(2,51)_ = 4.40, p=0.02). Notably, while morphine-treated females tested in adolescence required a greater number of trials and errors versus male morphine adolescent mice, no difference was observed in trials (p=0.836) or errors (p=0.697) to criterion in male versus female saline groups. In a more refined investigation into errors, we divided error type based on errors made prior to mice received less than six errors in one 16 trial bin (i.e. initial errors) and those after (regressive errors), the latter of which reflects an inability to maintain the rule strategy. No significant effect was observed on initial error types to criterion (sex: F(1,56)=0.33, p=0.57; treatment condition: F(2,56)=0.24, p=0.79; interaction: F(2,56)=0.01, p=0.99). Alternatively, morphine-treated females tested in adolescence exhibited greater regressive errors compared to saline-treated females, morphine-treated females tested in adulthood, and morphine-treated males tested in adolescence (interaction: F_(2,56)_=5.91, p=0.005).

**Figure 2.**
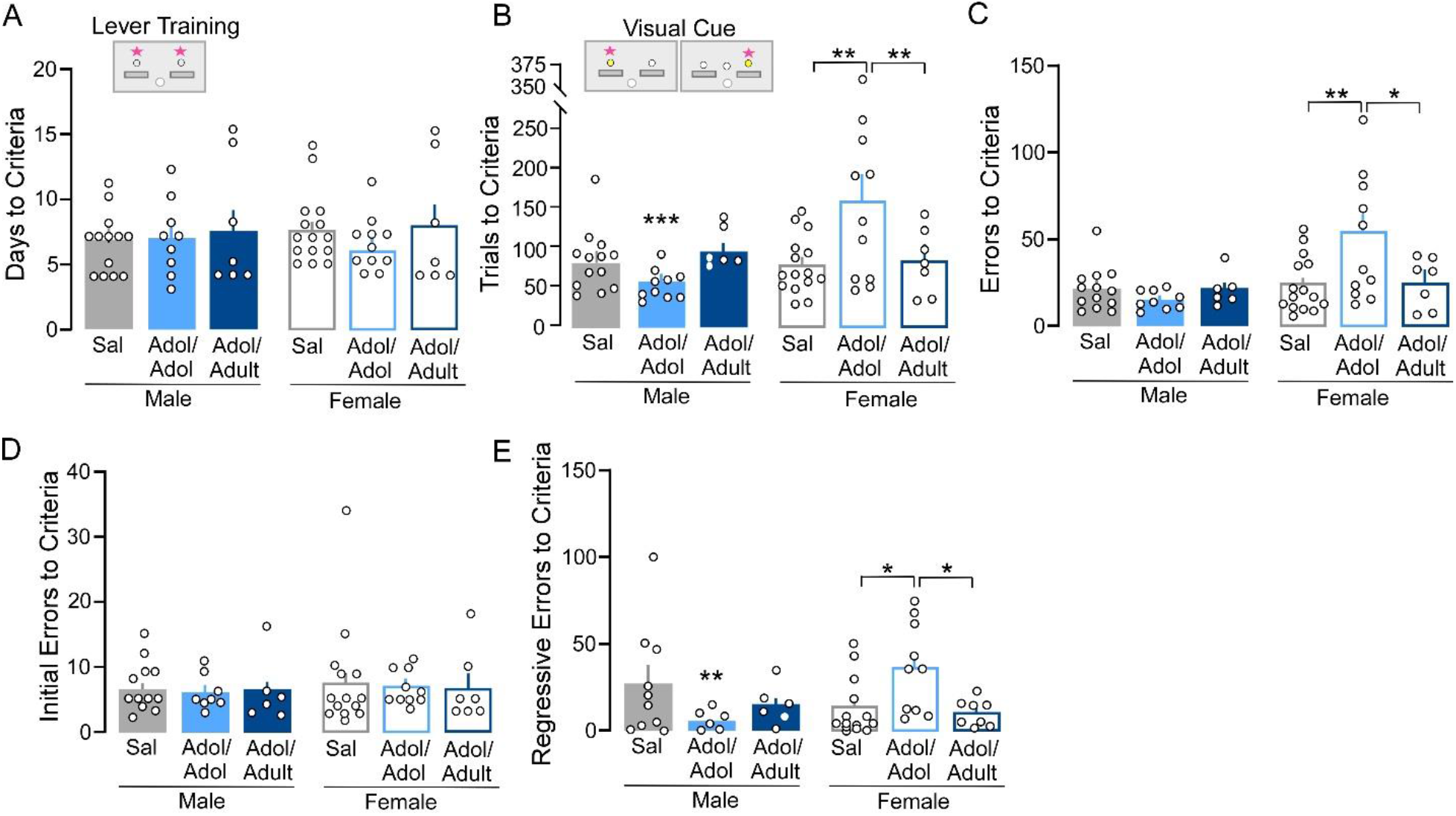
Lever training and visual cue test. **A)** No differences were observed in the number of days to criteria across treatment groups. **B-C)** Females treated with morphine and tested during adolescence require more trials **(B)** and exhibited more errors **(C)** to reach criteria during a visual cue test compared to females treated with saline, females treated with morphine but tested in adulthood, and males treated with morphine but tested in adolescence. **D-E)** No differences were observed across groups in the number of initial errors to criteria but females treated with morphine and tested in adolescence had more regressive errors compared to females treated with saline, females treated with morphine but tested in adulthood, and males treated with morphine but tested in adolescence. *p<0.05 **p<0.01,***p<0.001 vs. morphine-treated female mice tested in adolescence.

**Figure 3.**
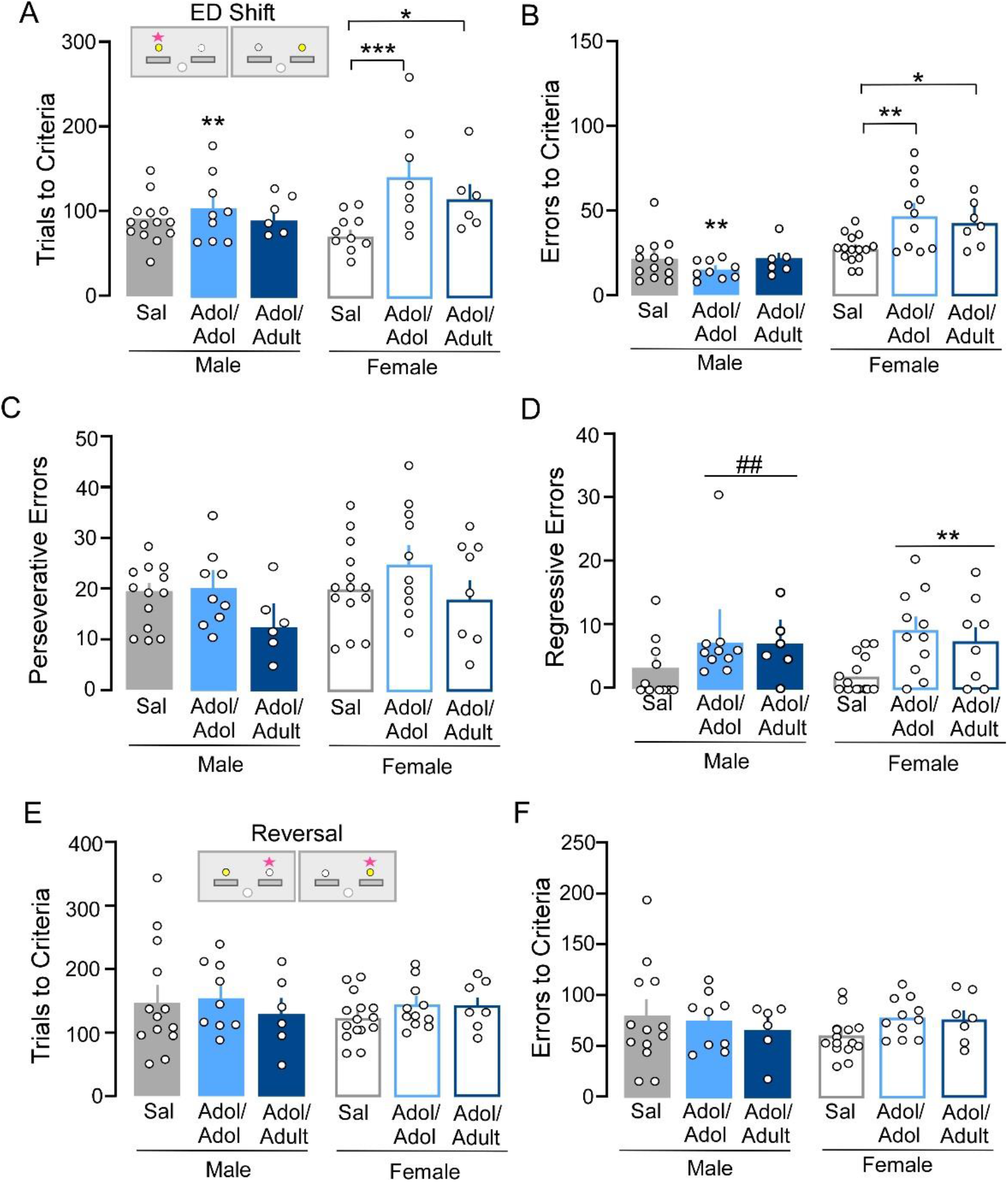
Extradimensional shift and reversal test. **A)** Trials to criteria and **B)** errors to criteria were greater in morphine-treated female mice tested in adolescence compared to saline-treated females, morphine-treated females tested in adulthoods, and morphine-treated males tested in adulthood. similar for all groups. **C)** The number of perseverative errors to criteria were similar across all treatment errors. D) Male and female mice treated with morphine had a greater number of regressive errors compared to their saline counterparts regardless of when testing occurred. E-F) The number of trials (E) and errors (F) to criteria were similar across all groups during a reversal test. *p<0.05, **p<0.01, ***p<0.001 vs. saline treated females. ^##^p<0.01 versus saline-treated males.

Comparison of performance during the extradimensional (ED) shift test showed an interaction of sex and treatment condition for number of trials (F_(2,61)_=3.82, P=0.028) and errors (F_(2,61)_=4.63, p=0.014) to reach criterion, with morphine treated females showing a greater number of both when tested during adolescence and adulthood compared to female saline controls. While morphine treated females tested in adolescence also showed a greater number of trials and errors to reach criterion compared to their morphine-treated male counterparts, no difference was observed between morphine females versus males tested in adulthood. In a more refined investigation into errors, we divided error type based on errors made prior to mice received less than six errors in one 16 trial bin (i.e. perseverative errors) and those after (regressive errors), the latter of which reflects an inability to maintain the rule strategy. Analysis of error types using a similar 16 trial bin as visual cue testing showed no effect of morphine on the number of perseverative errors to criterion (sex: F_(2,58)_=3.18, p=0.08; treatment condition: F_(2,58)_=2.92, p=0.06; interaction: F_(2,58)_=0.26, p=0.80). Alternatively, a main effect of treatment condition was observed for regressive type errors, with morphine treated mice exhibiting a greater number compared to saline controls regardless of testing age (sex: F_(3,58)_=0.52, p=0.48; treatment condition: F_(2,58)_=6.97., p=0.002; interaction: F_(2,58)_=0.48, p=0.62). During a subsequent reversal learning test was unaffected in both males and females (*trials to criterion*, sex: F(2,59)=0.05, p=0.82; treatment condition: F(2,59)=1.41, p=0.25; interaction: F(2,59)=0.25, p=0.78). Notably, no significant effects of treatment condition, sex, or interaction of sex x treatment condition were observed for omissions in the visual cue test, ED shift test, or reversal test (data not shown). These data, indicate that adolescent morphine exposure has a greater negative impact on female cognition compared to males and that these effects are unique depending on the cognitive construct and time of testing.

## Discussion

### Lack of effect of on affect- and motivation-related behavior

Negative affect following opioid exposure is known to occur throughout withdrawal and abstinence periods in human patients, partially triggering drug-seeking behaviors as a means of relief from these psychopathological states (Dole VP and Nyswander M, 1965;Wikler A, 1973). Further, in studies using male rats, anxiety-like behaviors were observed with precipitated withdrawal using the drug naloxone after repeated injection of morphine (Rothwell PE et al., 2010). There are many possibilities as to why no affect-related behavioral changes were observed in this experiment. One explanation is that the morphine doses used here were insufficient to promote physiological and/or psychological dependence compared to past studies which often use an increasing dose approach with final doses 10x higher (Madayag et al., 2019). While physical withdrawal symptoms were not assessed in the current study, morphine treatment did not significantly reduce body weight – a common phenomenon observed in preclinical dependence studies (Houshyar et al., 2003; Yanaura et al., 1975). Further, low dose self-administration of a fast-acting opioid has also previously been shown to not alter affect-related behavior. On the other hand, past studies observing changes in affective behavior were in rats and used a precipitated withdrawal model, whereas behavior was taken during spontaneous withdrawal in the present study. Finally, it is possible that our timeline of testing may have also impacted the detection of affective behavior changes, especially in mice tested during adulthood, 3-4 weeks following the cessation of drug exposure.

### Impact of adolescent morphine in cognition

Previous studies in adult mice have shown that intravenous self-administration of a low dose fast acting opioid impairs cognitive flexibility in males and females approximately 2-3-wk following drug exposure (Anderson et al., 2021). While these deficits were observed after 2-wk of self-administration in females, males required 4-wk of drug exposure for deficits to emerge. Here, we observed a similar female-specific deficit in flexibility that was present when tested in adolescence (∼2-wk following the final drug exposure) and persisted into adulthood (∼5-wk following drug exposure). Unlike adulthood, opioid exposure in the present study also impaired performance in a visual discriminative learning test, however these deficits did not persist into adulthood. Although cognitive flexibility was not tested during more protracted withdrawal, these findings indicate that adolescent morphine exposure may produce a more enduring deficit in flexibility when compared to exposure during adulthood. Further, it is possible that exposure during a critical development period for prefrontal cortices produces more widespread deficits in cognitive processing as performance in the discriminative learning task was also impaired. Notably, impaired performance in the visual cue test is not likely to reflect an overall deficit in associative learning processes, or changes in motivational states, as females do not take longer to acquire the initial lever training task. Further, it is unlikely that it reflects deficits in general attention to a cue or reduced visual acuity (Prusky GT et al., 2000), but rather the addition of a visual cue contingency not present during lever training is sufficient may act as a sufficient attentional shift (Anderson et al., 2021).

Previous studies with high-dose ethanol exposure have shown that females, but not males, are more susceptible to impairments in flexible behavior when exposed during adolescence, whereas the opposite pattern emerged following exposure during adulthood – highlighting an interaction between sex and age (Barker JM,Bryant KG,Osborne JI and Chandler LJ, 2017). While caution should be taken when comparing effects across different types of opioids and different exposure models (i.e., contingent versus noncontingent), it is possible that females remain more susceptible to deficits in cognitive processing compared to males regardless of age of exposure – highlighting a potential need to greater consideration when prescribing these medications.

